# An Event-Related Potential Study of Onset Primacy in Visual Change Detection

**DOI:** 10.1101/539932

**Authors:** Jennifer Van Pelt, Benjamin G. Lowe, Jonathan E. Robinson, Maria J. Donaldson, Patrick Johnston, Naohide Yamamoto

## Abstract

Onset primacy is a behavioural phenomenon whereby humans identify the appearance of an object (onset) with greater efficiency than other kinds of visual change, such as the disappearance of an object (offset). The default mode hypothesis explains this phenomenon by postulating that the attentional system is optimised for onset detection in its initial state. The present study extended this hypothesis by combining a change detection task and measurement of the P300 event-related potential, which was thought to index the amount of processing resources available to detecting onsets and offsets. In an experiment, while brain activity was monitored by electroencephalography, participants indicated the locations of onsets and offsets under the condition in which they occurred equally often in the same locations across trials. Although there was no reason to prioritise detecting one type of change over the other, onsets were detected more quickly and they evoked a larger P300 than offsets. These results suggest that processing resources are preferentially allocated to onset detection. This biased allocation may be a basis on which the attentional system defaults to the ‘onset detection’ mode.

---

The ability to shift the locus of attention within an environment provides an adaptive mechanism that is useful for the detection of change. In the absence of swift and successful change detection, it would be difficult to navigate everyday life. Driving a car or crossing the road, for example, requires the ability to detect new obstacles as they appear. Research involving visual search and detection paradigms has indicated that humans are adept at noticing new objects that abruptly enter their environment (Yantis & Jonides, 1984; for reviews, see Egeth & Yantis, 1997; Luck et al., 2021). This research showed that while there are other types of visual change that can be noticed well (e.g., object motion, colour alteration, and disappearance; Franconeri & Simons, 2003; Johnson et al., 2001; Pratt & McAuliffe, 2001), the abrupt appearance of a new object (object onset) tends to be particularly effective in capturing observers’ attention (Adams et al., 2023; Boot et al., 2005; Chua, 2013; Enns et al., 2001; Hillstrom & Yantis, 1994; Mounts, 2000). Under certain conditions, however, even significant changes in visual scenes go unnoticed, through a phenomenon known as change blindness (Simons & Rensik, 2005). Behavioural studies have demonstrated that the advantage of object onset as established in the visual search and detection paradigms is applicable to change blindness paradigms too; that is, object onset is more resistant to change blindness than changes in object colour, luminance, and motion (Cole & Liversedge, 2006; Cole et al., 2004) as well as the sudden disappearance of an object (object offset; Brockmole & Henderson, 2005; Cole & Kuhn, 2010; Cole et al., 2004). The comparative efficiency of onset over offset detection has been referred to in the literature as onset primacy (Cole et al., 2004; Donaldson & Yamamoto, 2012).

The persistence of onset primacy across experimental paradigms led Donaldson and Yamamoto (2016) to propose that onset detection is the default processing mode of the attentional system. To process other kinds of change, a shift is required from this default mode, resulting in less efficient responses. While the robust nature of onset primacy may be functionally adaptive, given that onset detection is advantageous in most situations, there are other situations in which offset detection may be more beneficial. For example, a lifeguard monitoring a crowded beach needs to notice the disappearing swimmer; a parent watching over a group of children in the playground needs to notice if one goes missing. As such, it is important to understand how and why onset primacy occurs by investigating the processes that underlie onset and offset detection.

In pursuing this endeavour, the present study tested the default mode hypothesis by examining differences in neural activation with electroencephalography (EEG) while participants attempted to detect onsets and offsets. According to the default mode hypothesis, the system of neural processing should initially be optimised for detection of onsets, making relevant areas of the brain more responsive to onsets than offsets. This enhanced processing of onsets should be reflected in event-related potentials (ERPs) that are of theoretical relevance to visual change detection. Specifically, this study focused on the P300 ERP as a neural marker of cognitive processes that underlie behavioural findings of onset primacy.

The P300, which is also called the P3, is a positive deflection that typically occurs between 300 and 500 ms after the onset of sensory stimuli (Hopfinger & Mangun, 1998; Hopfinger & Maxwell, 2005; Koivisto & Revonsuo, 2003), though it could range more widely from 250 ms up to 900 ms (Polich, 2007). It is generally implicated in information processing that involves selective attention and may be evoked after exposure to auditory or visual stimuli. Given that the P300 varies in topographic distribution, it is often conceptualised as two separate subcomponents, the P3a, a frontal distribution associated with stimulus novelty, and the P3b, a temporal-parietal distribution associated with attention and memory processing (Polich, 2007). The latter subcomponent has also been observed over the occipital areas in studies of selective attention (Koivisto & Revonsuo, 2003). While the P3a novelty subcomponent may instinctively be of interest in a change detection paradigm, it is likely that novelty effects would quickly be habituated across the repetitive presentation of visual stimuli, making it difficult to observe this subcomponent in averaged trials. The P3b subcomponent, however, remains of particular interest. As such, consequent discussion of the P300 is largely focused on the P3b in this paper. For simplicity, the P3b subcomponent is referred to as P300 hereafter.

A key reason for focusing on the P300 is that its amplitude is a good index of the amount of processing resources available for performing a task. This is best demonstrated in dual-task studies in which participants are given two tasks to perform simultaneously (Isreal et al., 1980; Mangun & Hillyard, 1990; Sirevaag et al., 1989; Wickens et al., 1983). A typical finding from these studies is that the P300 evoked by a primary task decreased when a secondary task was made more difficult so that it demanded a greater degree of participants’ attention (which, in turn, reduced their attention to the primary task). The P300 is modulated in the same way even when two tasks do not coincide strictly—there can be a delay of up to 1–1.5 s between them (Nash & Fernandez, 1996; Strayer & Kramer, 1990). These findings indicate that the P300 generally reflects the trade-off relationship between tasks when participants mentally prepare for performing both of the tasks, and its amplitude goes up and down as more and less resources are allocated to a task of interest. This idea is directly applicable to the current paradigm because there is similar reciprocity between onset and offset detection such that as observers improve their behavioural performance in detecting offsets through training, their efficiency in detecting onsets declines (Donaldson & Yamamoto, 2016). Taken together, it was postulated that P300 amplitude would function as a measure of processing resources allotted for detecting onsets and offsets.

The present study used this postulation to test the default mode hypothesis. This hypothesis posits that perceiving onsets is prioritised in the system of visual change detection in its initial state. One way of implementing this prioritisation is to assign a greater amount of processing resources for detecting onsets by default. Thus, if a larger amplitude of P300 was observed when participants detected onsets as compared to when they detected offsets, it would support the default mode hypothesis by showing that more resources are indeed allocated to detection of onsets. It is important to note that, as shown in the method section below, onsets and offsets were equal in the present experiment in that both were to-be-detected targets and they occurred in the same frequency and in the same spatial locations across trials. Therefore, participants should not have had any particular reasons to pay more attention to one type of change than the other. If onsets were still detected more efficiently, and if the detection was accompanied with a larger P300, it would indicate that allocation of processing resources is biased in favour of onset detection as the default mode of the system.

In summary, the goal of the present study was to seek neural evidence for the default mode hypothesis of onset primacy (Donaldson & Yamamoto, 2016). To this end, P300 amplitude was measured while participants attempted to detect onsets and offsets. It was predicted that the P300 would appear in a larger amplitude for onset than offset conditions in electrodes over temporal, parietal, and occipital regions of the brain.

## Method

The experiment reported below was approved by the Office of Research Ethics and Integrity of Queensland University of Technology (QUT) and conducted in accordance with the National Statement on Ethical Conduct in Human Research (National Health and Medical Research Council, 2018).

### Participants

Twenty-five QUT students (19 female, 6 male) aged 17–29 years (*M* = 20.88, *SD* = 3.18) participated in return for partial course credit. Written informed consent was obtained prior to their participation. All were right-handed with no known history of neurological disorder, and had normal or corrected-to-normal vision, as confirmed by a Snellen eye-chart.

### Materials

This experiment used the change detection task developed by Donaldson and Yamamoto (2012), who modelled it from Cole et al.’s (2003) one-shot flicker paradigm. Stimuli depicted visual scenes, each of which included a small circular table-top (38 cm diameter) with a single supporting leg (75 cm height). A range of objects of approximately equal size (4 × 3 × 2 cm of width, height, and depth) were placed onto the table in 16 different arrangements. The number of objects on the table was varied, ranging from six to nine, to minimise the potential to predict patterns of change. Images were presented at a central fixation point, with visual angle subtending approximately 7° horizontally and 4° vertically. Participants viewed the stimuli from an approximate distance of 60 cm on a monitor with a screen resolution of 1920 × 1080 pixels, via PsychoPy software (version 1.86; Peirce, 2007, 2009).

### Design and Procedure

Participants were informed that they were going to view a series of paired photographs, and that a change would be identifiable on the appearance of the second image. They were instructed to indicate whether they observed the change on the left or right side of the stimulus by pressing either F or J on their keyboard, respectively. Participants were asked to keep their index fingers resting on the response keys throughout the experiment and to respond as quickly and accurately as possible. Participants were not informed of the nature of change (i.e., onset or offset) and received no performance feedback.

In onset trials, the second image of a photograph pair contained one additional object on the tabletop; in offset trials, the second image removed one object from the tabletop (see Figure 1 for an example). The change occurred on each side of the table (left and right) an equal number of times and was counterbalanced across onset and offset conditions. To control for the potential influence of object properties, such as colour, location, or semantic salience, the same paired photographs were used for both onset and offset trials, in reversed order. Each photograph in the pair had either seven or eight objects on the tabletop, with each object acting as the change target an equal number of times. There were 64 trials for each change type, making a total of 128 experimental trials.

**Figure 1.**
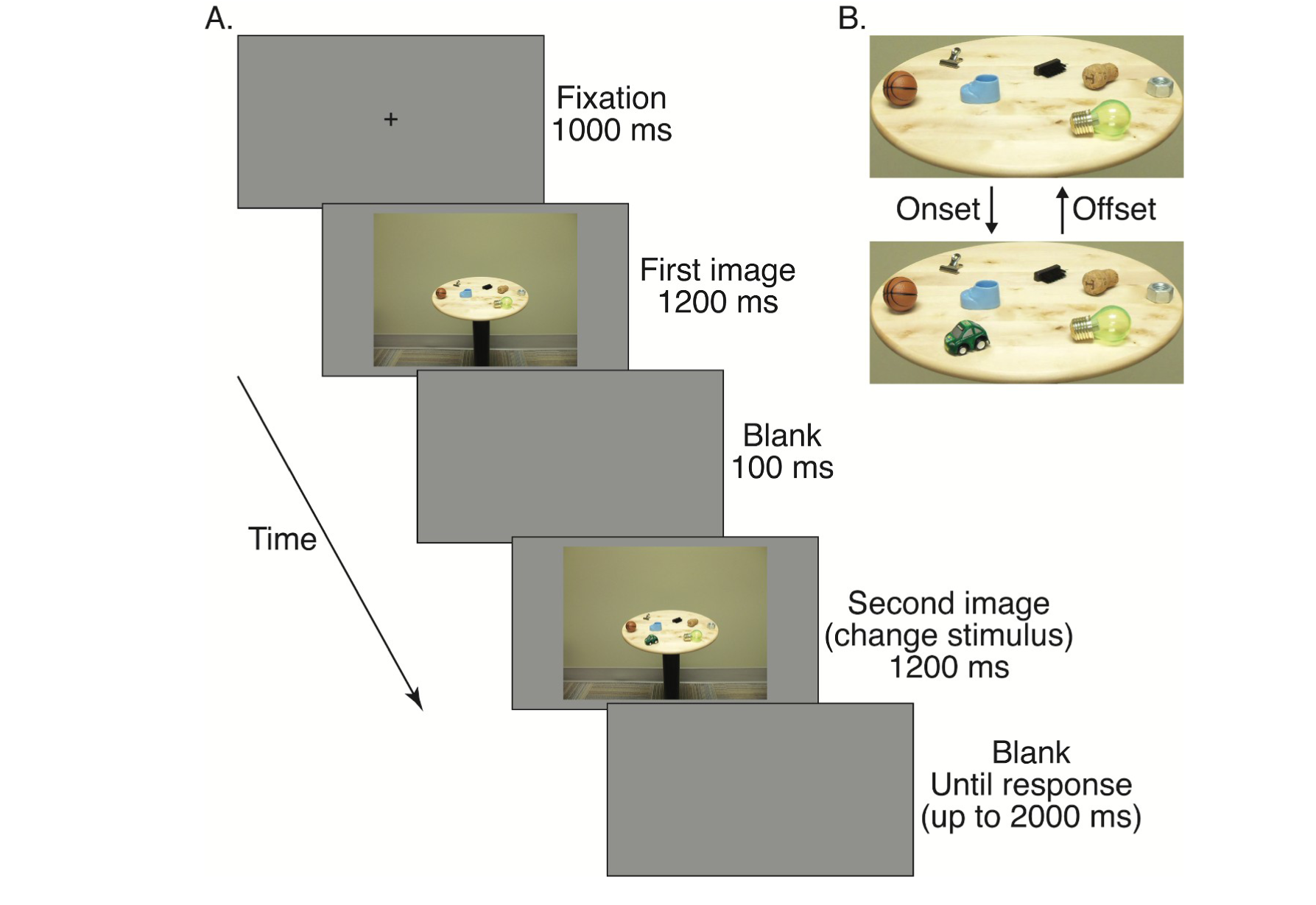
A Sample Experimental Trial. *Note.* (A) Sequence of trials. (B) A closer view of stimuli, demonstrating how the paired images changed in onset and offset trials. Here, the green car is the object of change. The figure is adapted from Donaldson and Yamamoto (2012).

Additional 32 photograph pairs were used for creating filler trials. In a filler onset trial, an eight-object display was followed by a nine-object display. In a filler offset trail, a seven-object display was followed by a six-object display. These filler trials (16 trials per change type) were included to discourage participants from anticipating the type of an upcoming trial on the basis of the first photograph alone. Without the filler trials, having a seven-object display in the first photograph could mean for certain that it would be an onset trial; similarly, having an eight-object display in the first photograph could mean that it would be an offset trial. The filler trials were intermixed with experimental trials, creating a total of 80 onset and 80 offset trials. These trials were randomly presented to each participant in a single block. Data from the filler trials were not included in the analysis.

Participants also completed eight onset and eight offset practice trials to familiarise themselves with the task. The practice trials were randomly presented in a block preceding the main block of experimental and filler trials. Photograph pairs used for the practice trials were not repeated in the main block.

The sequence of a trial is shown in Figure 1. First, participants viewed a central fixation cross for 1000 ms. The first image then appeared for 1200 ms, followed by a blank grey screen for 100 ms. The second image (referred to as *change stimulus* hereafter) then appeared for 1200 ms, before a final grey screen lasting 2000 ms. Participants were able to provide a response any time after the appearance of the change stimulus. When a response was provided, or when the final grey screen timed out, the fixation cross reappeared to indicate the start of the next trial.

### EEG Data Acquisition and Analysis

EEG data were recorded using the BioSemi ActiveTwo 64-channel amplifier and the ActiView software (version 7.06) at a sampling rate of 1024 Hz. The 64 EEG electrodes were laid out in accordance with the International 10–10 system, with a common mode sense and driven right leg circuit as online recording references. Electrode offsets were kept within ±25 mV prior to the recording.

Pre-processing was conducted using MNE-Python (version 1.4.2; Gramfort et al., 2013) within a Python 3.9.16 environment. It commenced with generating a second copy of a given participant’s raw EEG, which was high-pass filtered at 1 Hz to remove signal drifts so that independent components were identified (Winkler et al., 2015). These filtered copies of the data were produced solely for the purpose of identifying artefacts and were not used during any other subsequent analyses. Fifteen independent components were found through this process. The scalp topography and time series of these components were then visually inspected, and those that resembled eye-blink artefacts were removed from the original (i.e., unfiltered) data.

Subsequently, bandpass (0.1–30 Hz) and notch (50 Hz) filters were applied to the original EEG data, which were then downsampled to 1000 Hz. Next, the time series of each electrode was visually inspected, and when excessively noisy electrodes were found outside the regions of interest (ROIs; see Outcome Measures), they were interpolated via the spherical spline method (Perrin et al., 1989). On average, 3.48 interpolations were made per participant (*SD* = 3.36). Finally, data were re-referenced to the average activity of all 64 electrodes.

Data were segmented into epochs between −100 and 700 ms relative to the appearance of each image and baseline-corrected to the mean of a pre-stimulus period (−100– 0 ms). Epochs containing filler trials or incorrectly performed experimental trials were excluded from analysis. Out of the remaining epochs, those in which peak-to-peak difference between −100 and 600 ms exceeded 200 μV in any of the electrodes included in the ROIs were further removed. For three participants, no epochs survived these processes in one or more conditions, resulting in exclusion of these participants from all analyses. Participant-level ERPs were then averaged across epochs per condition.

### Outcome Measures

#### Behavioural Measures

Reaction time was measured as the time that elapsed between the appearance of a change stimulus and a participant’s response. Accuracy of the change location judgement, as indicated by the keyboard button press, was also measured. Trials in which participants failed to provide a response before the end of the final grey screen were considered incorrect. All trials (i.e., including those that were rejected for EEG analysis due to recording artefacts) were used for reaction time and accuracy calculations, with the exception that incorrectly performed trials were excluded from reaction time analysis.

#### P300

The ROIs for the P300 included electrodes over the cortical surface of the temporal, parietal, and occipital lobes (Eimer & Mazza, 2005; Koivisto & Revonsuo, 2003; Polich, 2007). They were grouped into clusters corresponding to their topographic location (Figure 2): left (CP1, CP3, CP5, TP7, P1, P3, P5, P7, PO3, PO7, and O1), centre (CPz, Pz, POz, and Oz), and right (CP2, CP4, CP6, TP8, P2, P4, P6, P8, PO4, PO8, and O2). The mean amplitude measures at these electrode sites were computed by temporally averaging amplitude values across time points within the time window of 275–500 ms after the appearance of a change stimulus, which was set a priori by referring to time windows used in previous studies for measuring the P300 (Hopfinger & Mangun, 1998; Hopfinger & Maxwell, 2005; Koivisto & Revonsuo, 2003). The mean amplitude was calculated for each electrode first, and then averaged across the electrodes per ROI before being entered into statistical models for analysis.

**Figure 2.**
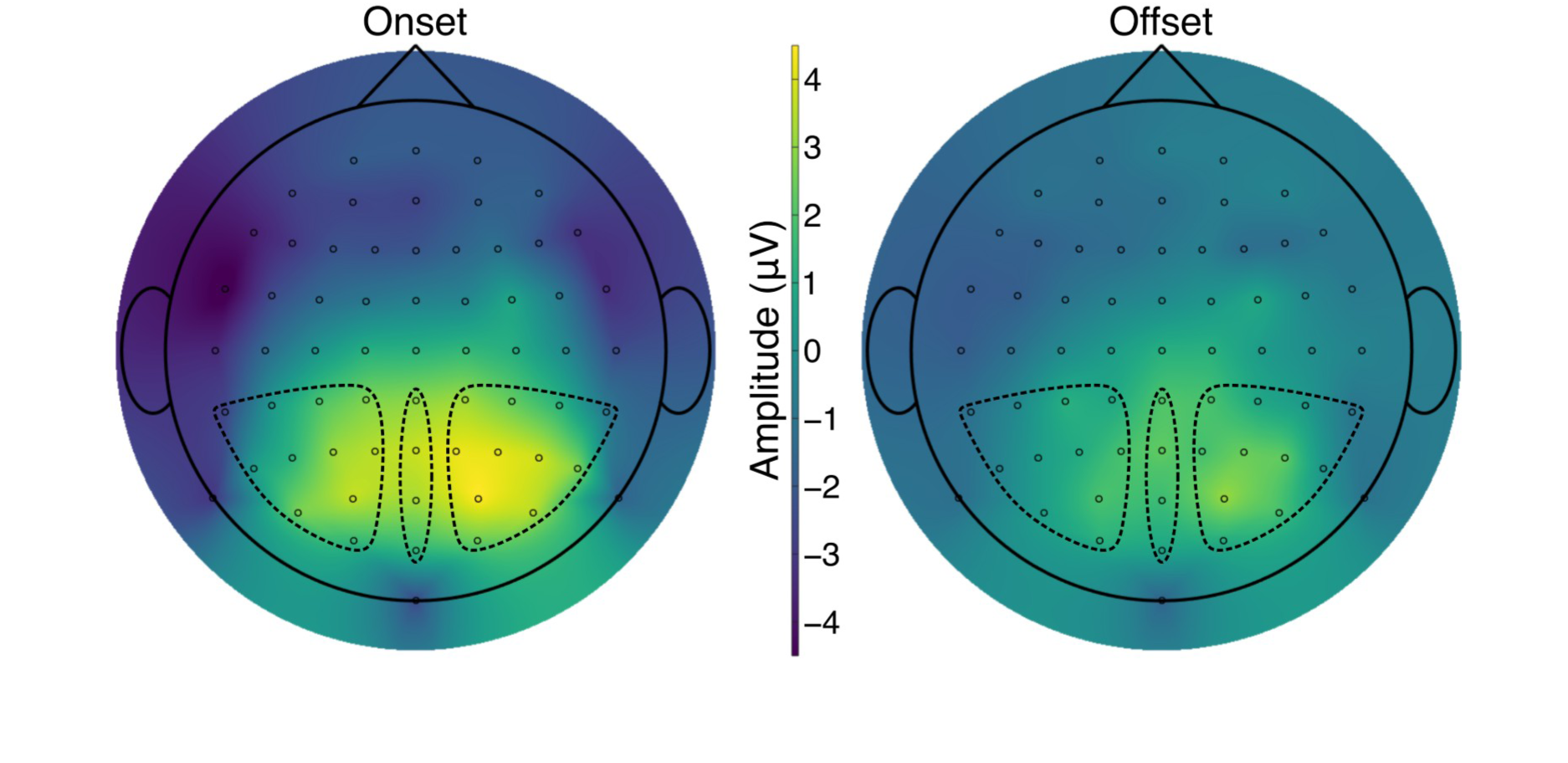
Grand-Average Topographic Plots in the P300 Time Window. *Note*. The plots show the distributions of the mean voltage in the period of 275–500 ms post change stimulus averaged across participants separately for each change type. Circles represent EEG electrodes. Broken lines indicate electrode clusters that constituted the regions of interest (ROIs): left (CP1, CP3, CP5, TP7, P1, P3, P5, P7, PO3, PO7, and O1), centre (CPz, Pz, POz, and Oz), and right (CP2, CP4, CP6, TP8, P2, P4, P6, P8, PO4, PO8, and O2).

Peak amplitude measures were also derived by finding the largest positive amplitude peaks within the same time window for individual electrodes and averaging them per ROI. The mean and peak amplitude data yielded consistent results. Thus, for brevity, analysis of the peak amplitude data is not reported below. Both mean and peak amplitude values at each electrode site are available on the Open Science Framework at https://osf.io/jnq27.

## Results

### Behavioural Results

#### Accuracy

For each type of change, accuracy scores were defined as outliers, if they were more than two standard deviations away from the mean of 22 participants (i.e., all participants except those who were excluded due to EEG recording artefacts). Two scores in the onset condition (73.44% and 78.13%) and two scores in the offset condition (70.31% and 71.88%), which were produced by three participants, met the criterion. These participants were further removed from all subsequent analyses.^1^ The mean accuracy scores of the remaining 19 participants (16 female, 3 male, 17–29 years of age with *M* = 20.89, *SD* = 3.19) were 96.13% (*SD* = 3.22%) in onset trials and 94.65% (*SD* = 3.01%) in offset trials, which were statistically indistinguishable from each other, *t*(18) = 1.43, *p* = .171, *d_rm_* = 0.46, 95% CI [−0.70, 3.66] (for the definition of *d_rm_*, see Lakens, 2013). This result is not contrary to prediction, as previous studies have demonstrated that accuracy is not as sensitive to onset primacy as reaction time (Donaldson & Yamamoto, 2012, 2016). The similarly high accuracy in onset and offset detection ensured that analysis of EEG data reported below was performed on equivalent numbers of trials in onset and offset conditions.

#### Reaction Time

The mean reaction times across trials were 563.74 ms (*SD* = 72.49 ms) in the onset condition and 627.54 ms (*SD* = 50.24 ms) in the offset condition, indicating that participants detected onsets faster than offsets, *t*(18) = 4.82, *p* < .001, *d_rm_* = 0.94, 95% CI [35.99, 91.60]. These data align with prediction and show that onset primacy was present.

#### Relationship between Accuracy and Reaction Time

To ensure that the quicker detection of onsets than offsets was not due to speed-accuracy trade-offs, the relationship between accuracy scores and reaction times was examined at the level of individual participants. Specifically, for each participant, the mean accuracy score of the offset condition was subtracted from that of the onset condition, and the mean reaction time of the onset condition was subtracted from that of the offset condition. If these two measures negatively correlated across the participants, it could suggest that the participants sped up in onset trials by performing them less carefully. However, the correlation was positive, *r*(17) = .52, *p* = .024, 95% CI [.08, .79], showing that those who had greater degrees of advantage in the speed of onset detection tended to exhibit larger magnitudes of onset primacy in accuracy as well. This pattern is contrary to what would have been produced by speed-accuracy trade-offs, supporting the conclusion that onset primacy was robustly demonstrated by the behavioural data.

### EEG Results

For the 19 participants whose ERPs were assessed, on average, 90.58% of epochs (*SD* = 8.92%) remained in the analysed data after removal of those that were compromised by recording artefacts and incorrect behavioural responses. The mean P300 amplitude values were examined by a 2 (onset and offset) × 3 (left, centre, and right ROIs) repeated-measures analysis of variance (ANOVA).^2^ There was no evidence for violation of sphericity as per Mauchly’s tests (*W*s > .89, *p*s > .394).

Descriptive statistics for the mean amplitude data are displayed in Table 1. Consistent with prediction, the P300 amplitude was higher in onset than offset conditions. The amplitude values also suggest that amplitude varied with electrode location, with the amplitude being larger in the central and right ROIs than in the left ROI on average. These patterns can be seen in Figure 2 that shows the topographic distributions of the P300 amplitude. Figure 3 displays mean ERP waveforms from each ROI that demonstrate the expected pattern of the P300, wherein the amplitude was greater in onset trials than in offset trials.

**Figure 3.**
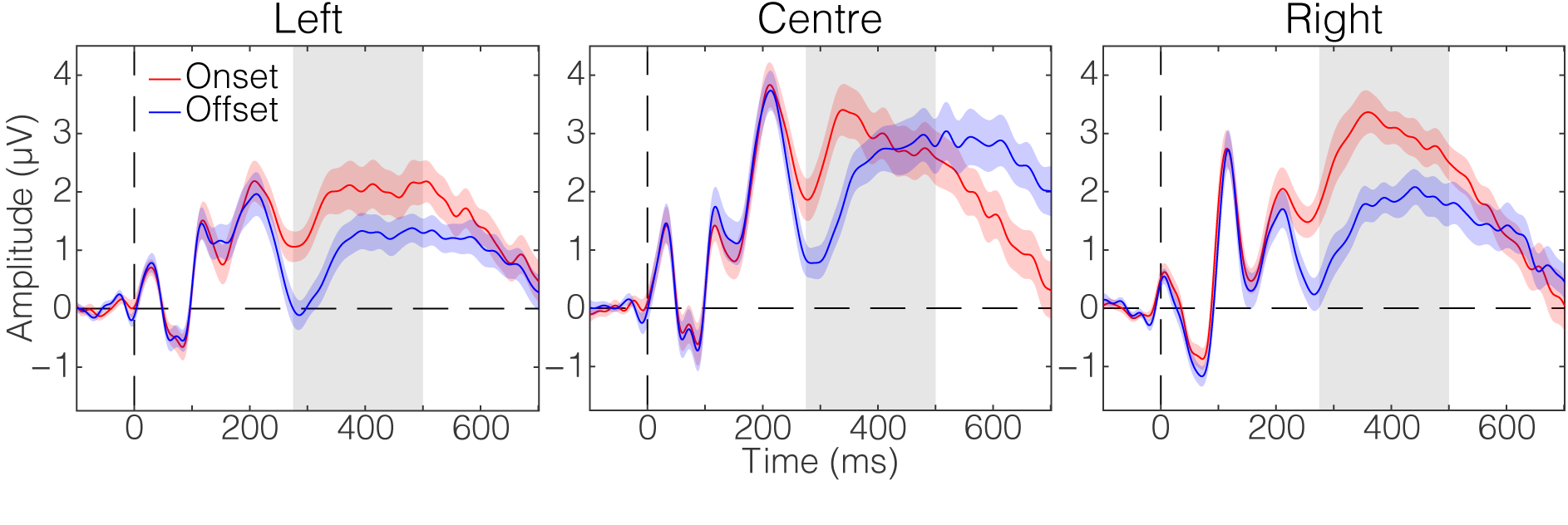
P300 Amplitude Time Series. *Note*. These waveforms were derived by averaging across participants, separately for each change type and each region of interest (left, centre, and right electrode clusters). Shading around each waveform represents ±1 standard error of the mean at each time point. Grey shaded areas indicate the time window from which P300 amplitude values were calculated for analysis (275–500 ms post change stimulus).

**Table 1.** Mean Amplitude of the P300.

| ROI | Onset |  |  | Offset |  |  |
| --- | --- | --- | --- | --- | --- | --- |
|  | *M* | *SD* | 95% CI | *M* | *SD* | 95% CI |
| Left | 1.87 | 1.70 | [1.02, 2.71] | 0.95 | 1.15 | [0.38, 1.52] |
| Centre | 2.83 | 1.90 | [1.89, 3.77] | 2.20 | 1.64 | [1.39, 3.01] |
| Right | 2.90 | 1.46 | [2.17, 3.62] | 1.61 | 1.26 | [0.99, 2.24] |

*Note.* The amplitude values are shown in $\mu\text{V}$ . ROI = region of interest.

The ANOVA examining the amplitude data revealed a significant main effect of change types, *F*(1, 18) = 21.11, *p* < .001, η*_G_*^2^ = .086, in which onsets (*M* = 2.53 µV, *SD* = 1.51 µV) were greater than offsets (*M* = 1.59 µV, *SD* = 1.05 µV). There was also a significant main effect of ROIs, *F*(2, 36) = 6.87, *p* = .003, η_G_ ^2^ = .086. Follow-up pairwise comparisons (each with a Bonferroni-corrected α of .016) indicated that this main effect emerged because EEG activity was lateralised towards the right hemisphere: While the mean amplitude values in the central (*M* = 2.51 μV, *SD* = 1.66 μV) and right (*M* = 2.25 μV, *SD* = 1.31 μV) ROIs were statistically indistinguishable from each other, *t*(18) = 0.84, *p* = .415, *d_rm_* = 0.17, 95% CI [−0.40, 0.92], the mean amplitude value in the left ROI (*M* = 1.41 μV, *SD* = 1.32 μV) was significantly lower than that in the central ROI, *t*(18) = 3.77, *p* = .001, *d_rm_* = 0.70, 95% CI [0.49, 1.72]. The difference between the left and right ROIs did not reach statistical significance, *t*(18) = 2.57, *p* = .019, *d_rm_* = 0.62, 95% CI [0.15, 1.53]. The interaction between change types and ROIs was not significant, *F*(2, 36) = 2.97, *p* = .064, η_G_ ^2^ = .007, suggesting that the way the effects of change types occurred did not substantially differ between the electrode clusters.

One possible confound in this experiment was that change stimuli in onset trials contained eight objects, whereas those in offset trials displayed seven objects. Thus, any effects of change types could have been caused by this simple difference in the number of objects. To empirically rule out this possibility, EEG responses to the first photograph of each pair were analysed. As shown in Supplementary Material, there was no evidence for P300 difference between onset and offset trials in this analysis, suggesting that the patterns of P300 amplitude evoked by the change stimuli were not mere artefacts of the experimental paradigm.

## Discussion

The current study aimed to test the default mode hypothesis (Donaldson & Yamamoto, 2016) by examining whether behavioural findings of onset primacy were reflected in EEG-recorded neural activation during a change detection task. The default mode hypothesis postulates that a larger amount of processing resources is allocated to onset detection than offset detection under the initial mode of attention, leading unbiased observers to perform trials involving the detection of onsets with greater neural efficiency than trials involving the detection of offsets. On the basis of previous studies of dual-task performance (Isreal et al., 1980; Mangun & Hillyard, 1990; Nash & Fernandez, 1996; Sirevaag et al., 1989; Strayer & Kramer, 1990; Wickens et al., 1983), the amplitude of the P300 ERP was hypothesised to be an index of this efficiency. Specifically, it was predicted that the P300 would have a higher amplitude in onset trials than in offset trails, across the specified time-window (275–500 ms post change stimulus) and ROIs (temporal, parietal, and occipital regions), reflecting the relative abundance of processing resources for detecting onsets in the change detection task.

The prediction was confirmed, as mean amplitude of the P300 was larger in onset than offset conditions, and this pattern emerged while participants detected onsets more quickly than offsets. Accuracy of change detection did not statistically differ between onset and offset trials, indicating that the quicker detection of onsets was not a mere consequence of speed-accuracy trade-offs. It should be noted that these results were obtained when the onset and offset trials were well equated—that is, onsets and offsets occurred equally often on the same objects and at the same locations across the trials, and the participants were instructed to respond to the location of a change, not to the type of the change. Given the design of the task, it was very likely that the participants carried out the trials without overtly shifting their attentional priority to specifically detecting onsets or offsets. Nevertheless, onsets were still detected faster and this behavioural performance was associated with the greater P300 amplitude. Together, these findings support the interpretation that onset primacy is a result of the attentional system’s default mode in which the allocation of processing resources is biased in favour of onset detection.

In addition to specifically supporting the default mode hypothesis, the present results are more broadly consistent with previous studies on change blindness and detection. Using tasks in which changes were difficult to perceive, these studies found that the P300 was evoked following successful detection of change (Koivisto & Revonsuo, 2003; Niedeggen et al., 2001). In the current study, it appears that not only onsets, but also offsets, elicited a canonical P300 component when their location was correctly indicated (Figure 3). Because the location judgements were made with very high accuracy, this study alone does not clarify whether the P300 is uniquely associated with correct recognition of change—that is, no reliable EEG data were available as to whether the P300 was absent when the changes were incorrectly localised or entirely missed. However, combined with the previous studies, these findings suggest that the P300 reflects neural processes that have to do with establishing conscious awareness of a visual stimulus and making decisions about the detected stimulus (Eimer & Mazza, 2005; Turatto et al., 2002).

Although the current study was carried out without forming any predictions about the effect of electrode location, differences in amplitude did emerge as a function of electrode location for the P300 component. Specifically, the amplitude was overall higher across electrodes in the midline and in the right hemisphere than those in the left hemisphere, regardless of change type. Moreover, there was a distinct peak in the right hemisphere in the onset condition (Figure 2). These amplitude patterns are consistent with results from previous studies in which the P300 was evoked in midline electrodes by appearing and disappearing visual targets (Eimer & Mazza, 2005; Hopfinger & Mangun, 1998, 2001), and also with the common view that the right hemisphere tends to be dominant in tasks involving visuospatial attention (Corballis, 2003; Mesulam, 1999; Shulman et al., 2010). This account is supported by EEG studies that examined performance across visuospatial tasks and identified similar P300 lateralisation (Alexander et al., 1995; Makeig et al., 1999).

A challenge in elaborating on the current P300 results is that this ERP component can be observed in multiple paradigms (Polich, 2007). Notably, the P300 is elicited when observers perceive rare and deviant stimuli within a sequence that mainly consists of frequent and standard stimuli (i.e., the oddball paradigm; Picton, 1992). At a glance, the oddball P300 can seem inapplicable to this study in which participants performed the same number of onset and offset trials. One way of reconciling the findings from the present and oddball paradigms is to conceptualise the P300 as the brain’s response to stimuli of potential significance (Ferrari et al., 2010)—both newly appearing objects and rarely occurring non-standard stimuli are worth perceiving because they may signify the occurrence of novel situations that require close attention. Consistent with this view, it has been shown that onsets and oddballs do not evoke the P300 when they are task-irrelevant and viewed passively (Bennington & Polich, 1999; Hopfinger & Maxwell, 2005). When interpreted in this framework, the enhanced P300 observed in the onset trials offers empirical support to the conceptual idea about why onset primacy occurs: It is often explained by theoretically assuming that onsets should call for prioritised processing because without having a conscious understanding of what the new objects are, it is not possible to react to them appropriately (and this reaction may need to be made quickly, like when the onsets pose a threat); on the other hand, offsets do not always demand such preferential processing because the disappearing objects have once been perceived (Cole et al., 2003; Donaldson & Yamamoto, 2016). The P300 amplitude difference between onset and offset conditions suggests that the neural system is in fact designed to assign greater significance to onsets than offsets, heightening its readiness for onset perception via biased deployment of attentional resources.

## Future Directions

Finally, it may be worth noting what steps can be taken to advance this research. Since this was the first study in which Donaldson and Yamamoto’s (2012) onset primacy paradigm was combined with EEG, participants’ reaction times were measured simultaneously with EEG recordings. This was to ensure that any patterns of EEG data were observed while onset primacy was actually taking place. Without reaction time data, it would have been necessary to assume that participants were detecting onsets more efficiently than offsets, but it is an open question whether onset primacy occurs in the same way when no speeded responses to the changes are required. Thus, in this initial investigation, there was a clear benefit of obtaining the reaction time data. However, it inevitably came with some drawbacks. Most notably, the EEG data reflected not just perceptual and cognitive processes inherent in onset primacy but also motor processes involved in response preparation and execution. Presumably, these motor processes were commonly engaged in onset and offset trials, and therefore their effects should not have caused a fundamental problem in the comparison between the two trial types. Nevertheless, to isolate EEG signals that are unique to onset primacy itself, it would be useful to conduct experiments in which participants do not make any behavioural responses while they view onset and offset stimuli. For example, they may be asked to respond to the stimuli upon receiving delayed cues. To prevent them from preparing specific responses during the viewing and delay periods, tasks may be designed such that response options vary trial-by-trial and only the cues specify which option is relevant in each trial. Such delayed-response paradigms are now justified because the present study has provided a clear EEG marker of onset primacy that future experiments can look for (i.e., the P300). These experiments would offer further clarification of how onsets and offsets modulate the EEG signals, thereby delineating onset primacy at both cognitive and neural levels.

Another way of expanding the present study is to compare object onset not just with object offset but with other kinds of visual change. The present experiment focused on onset and offset detection so as to make a tight comparison between these change types. They were created from the same images by reversing the order of presentation, making onset and offset trials equivalent in terms of stimulus properties such as object size, location, colour, and luminance. This manipulation was crucial because any one of these properties could cause a confound and obscure whether the enhancement of the P300 component should be attributed to object onset. However, to build on the rich behavioural literature in which onset primacy has been investigated by contrasting onsets with a wide range of visual events (e.g., changes in object colour, luminance, and motion; Adams et al., 2023; Cole & Liversedge, 2006; Cole et al., 2004; Enns et al., 2001; Franconeri & Simons, 2003; Hillstrom & Yantis, 1994; Johnson et al., 2001; Mounts, 2000), it is important to examine whether the amplitude of the P300 continues to be increased in the onset condition when it is compared against these non-offset conditions. Such research will more firmly establish the notion that the P300 is an EEG marker of onset primacy in visual change detection.

## Acknowledgements

The authors thank Yasmin Allen-Davidian for cooperation in participant recruitment.

## Declarations

### Ethics approval

This research was approved by the QUT Office of Research Ethics and Integrity (approval numbers 1700000238 and 5663).

### Consent to participate

Written informed consent was obtained from participants.

### Consent for publication

Participants consented to publish non-identifiable data.

### Availability of data and materials

Data collected and analysed in this research are available at https://osf.io/jnq27.

### Code availability

Code used for processing and visualising EEG results is available at https://osf.io/jnq27.

### Authors’ contributions

JVP: conceptualisation, data curation, formal analysis, investigation, methodology, visualisation, writing—original draft; BGL: data curation, formal analysis, software, validation, visualisation, writing—review & editing; JER: data curation, formal analysis, methodology, software, writing—review & editing; MJD: conceptualisation, methodology, resources, writing—review & editing; PJ: formal analysis, methodology, resources, software, supervision, writing—review & editing; NY: conceptualisation, data curation, formal analysis, funding acquisition, methodology, project administration, resources, software, supervision, visualisation, writing—review & editing

### Open Practices Statement

The availability of research materials is as declared above. The experiment was not pre-registered.

## Footnotes

1 Both behavioural and EEG data were analysed by including these participants too, and results remained unchanged.

2 A 2 × 3 × 2 ANOVA was also run by including the location of change (left and right sides) as an additional within-participant factor. Given that fixation and eye movements were not strictly controlled in this experiment, it was expected that the location of change would not show any major effects. Consistent with this expectation, this factor yielded no significant main effect or interactions.

