## Supplementary Material for "An Event-Related Potential Study of Onset Primacy in Visual Change Detection"

### Analysis of P300 Amplitude Evoked by the First Image of Each Photograph Pair

The main analysis reported in the article revealed that change stimuli (the second images of photograph pairs) evoked greater P300 amplitude in onset trials than in offset trials. One potential issue in this analysis was that onset and offset conditions differed in the number of objects they used in the change stimuli (onset: eight objects; offset: seven objects). To rule out the possibility that the larger P300 amplitude in the onset condition was a mere consequence of having one more object in the change stimuli, electroencephalographic (EEG) responses to the first images of the photograph pairs were analysed in the same way as in the main analysis. In this supplementary analysis, the onset trials showed seven objects, whereas the offset trials displayed eight objects. As in the main analysis, both mean and peak P300 amplitude values were examined. Since they led to the same conclusion, only the mean amplitude values are reported below for brevity. The mean and peak amplitude data from the supplementary analysis are available on the Open Science Framework at https://osf.io/jnq27.

Figure S1 plots topographic distributions of mean amplitude values in the P300 time window (275–500 ms after the appearance of first images). Figure S2 displays mean waveforms from each region of interest (ROI). Table S1 summarises descriptive statistics for the mean P300 amplitude data. They all indicate that onset and offset trials yielded very similar patterns of the P300.

Mean P300 amplitude values were examined by a 2 (onset and offset) × 3 (left, centre, and right ROIs) repeated-measures analysis of variance. Neither the main effect of change types nor the interaction between change types and ROIs was significant, $F(1, 18) = 0.25$, $p = .624$, $\eta_G^2 < .001$ and $F(2, 36) = 1.21$, $p = .311$, $\eta_G^2 = .002$, respectively. The main effect of ROIs was significant, $F(2, 36) = 3.92$, $p = .029$, $\eta_G^2 = .047$, suggesting that EEG activity was lateralised towards the right hemisphere (left ROI: $M = 3.05$ μV, $SD = 1.27$ μV; central ROI: $M = 2.92$ μV, $SD = 2.08$ μV; right ROI: $M = 3.87$ μV, $SD = 2.02$ μV). However, none of the follow-up pairwise comparisons (each with a Bonferroni-corrected α of .016) reached statistical significance, $t$s(18) < 2.39, $p$s > .028, $d_{rm}$s < 0.46 (for the definition of $d_{rm}$, see Lakens, 2013).

In sum, the supplementary analysis did not yield any evidence that onset and offset conditions were different in P300 amplitude, even though they still differed in the number of objects they displayed (in this case, offset trials showed one more object than onset trials did). These results support the interpretation that the difference in P300 amplitude evoked by change stimuli mainly stemmed from detection of onsets versus offsets, rather than merely reflecting the fact that the change stimuli in the onset condition contained an extra object as compared with those in the offset condition.

**Figure S1**

*Grand-Average Topographic Plots in the P300 Time Window for the Supplementary Analysis*

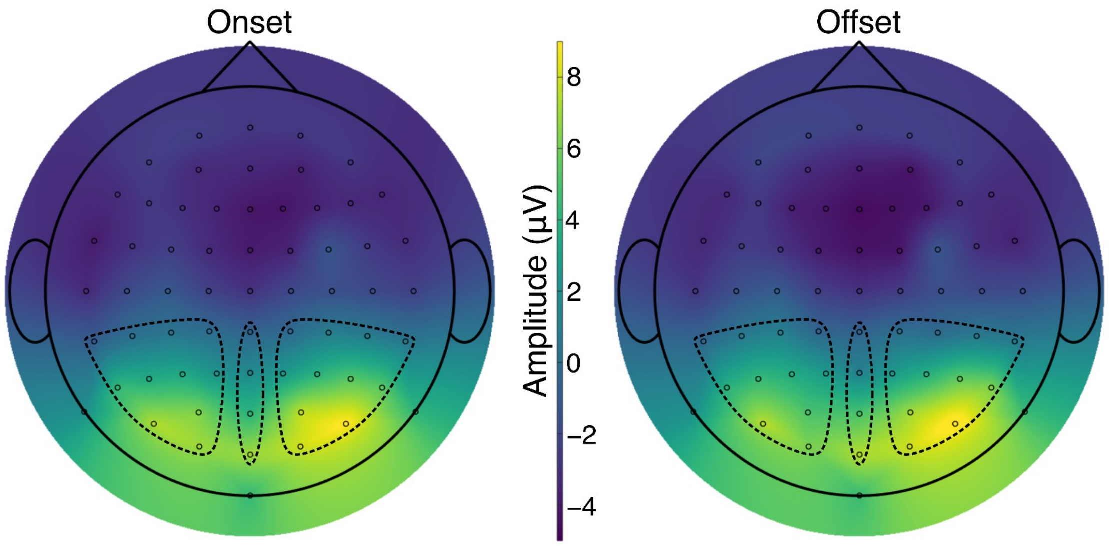


*Note*. The plots show the distributions of the mean voltage in the period of 275–500 ms after the appearance of first images, averaged across participants separately for each change type. Circles represent EEG electrodes. Broken lines indicate electrode clusters that constituted the regions of interest (ROIs): left (CP1, CP3, CP5, TP7, P1, P3, P5, P7, PO3, PO7, and O1), centre (CPz, Pz, POz, and Oz), and right (CP2, CP4, CP6, TP8, P2, P4, P6, P8, PO4, PO8, and O2).

**Figure S2**

*P300 Amplitude Time Series* for the Supplementary Analysis

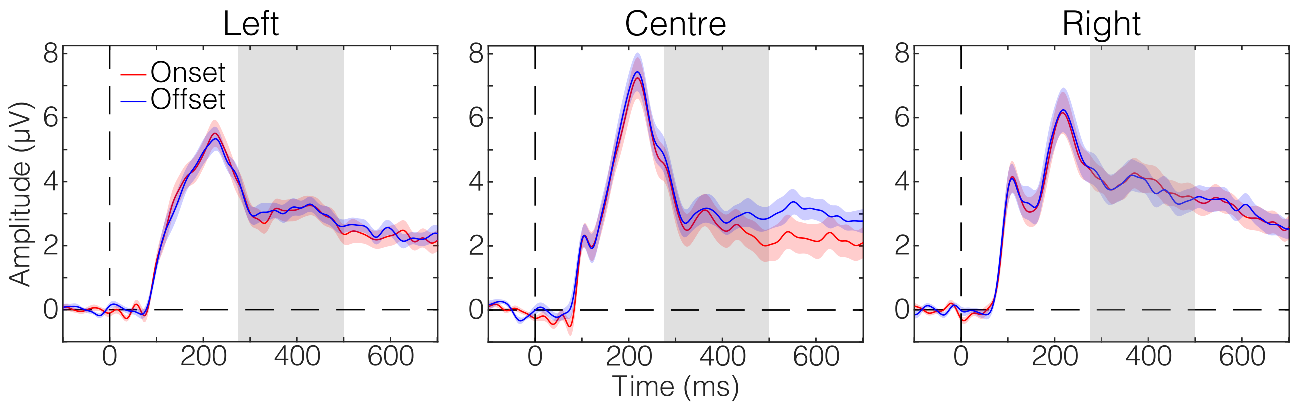


*Note*. These waveforms were derived by averaging across participants, separately for each change type and each region of interest (left, centre, and right electrode clusters). Shading around each waveform represents ±1 standard error of the mean at each time point. Grey shaded areas indicate the time window from which P300 amplitude values were calculated for analysis (275–500 ms after the appearance of first images).

**Table S1**

*Mean Amplitude of the P300 in the Supplementary Analysis*

| ROI | Onset | | | Offset | | |
|---|---|---|---|---|---|---|
| | *M* | *SD* | 95% CI | *M* | *SD* | 95% CI |
| Left | 3.03 | 1.52 | [2.28, 3.79] | 3.07 | 1.16 | [2.50, 3.65] |
| Centre | 2.75 | 2.39 | [1.57, 3.93] | 3.08 | 1.93 | [2.12, 4.04] |
| Right | 3.92 | 2.06 | [2.89, 4.94] | 3.83 | 2.09 | [2.79, 4.86] |

*Note*. The amplitude values are shown in μV. ROI = region of interest.
